# Gaze-contingent processing improves mobility performance and visual orientation in simulated head-steered prosthetic vision

**DOI:** 10.1101/2023.09.18.558225

**Authors:** Jaap de Ruyter van Steveninck, Mo Nipshagen, Marcel van Gerven, Umut Güçlü, Yağmur Güçlüturk, Richard van Wezel

## Abstract

The enabling technology of visual prosthetics for the blind is making rapid progress. However, there are still uncertainties regarding the functional outcomes, which can depend on many design choices in the development. In visual prostheses with a head-mounted camera, a particularly challenging question is how to deal with the gaze-locked visual percept associated with spatial updating conflicts in the brain. A recently proposed compensation strategy is gaze-contingent image processing with eyetracking, which enables natural visual scanning and reestablished spatial updating based on eye movements. The current study evaluates the benefits of gaze-contingent processing versus gaze-locked and gaze-ignored simulations in the context of mobility and orientation, using a simulated prosthetic vision paradigm with sighted subjects. Compared to gaze-locked vision, gaze-contingent processing was found to improve the speed in all experimental tasks, as well as the subjective quality of vision. Similar or further improvements were found in a control condition that ignores gaze-depended effects, a simulation that is unattainable in the clinical reality. Our results suggest that gaze-locked vision and spatial updating conflicts can be debilitating for complex visually-guided activities of daily living such as mobility and orientation. Therefore, for prospective users of head-steered prostheses with an unimpaired oculomotor system, the inclusion of a compensatory eye-tracking system is strongly endorsed.

## 1. Introduction

Worldwide, approximately 40 million people are affected by blindness (1). In the past few decades promising progress has been made towards the development of visual prosthetics for the blind that aim to support basic visually-guided activities of daily living, increasing the users’ autonomy and overall quality of life (2–6). The technology of prosthetic vision is still in the early stages of development, and currently the clinical potential is explored for various designs. Even given the rapid recent developments (for instance, see (7, 8)) it is clear that the quality of the prosthetic vision will be relatively elementary compared to natural vision. This forms the motivation for an active line of research studying the expected functionality and utility in relation to specific design choices in the development (9, 10).

A particularly challenging question is how visual prosthetics will deal with the intricate neural interactions between visual processing and oculomotor behavior. In the human brain, eye movements are continuously monitored to preserve visual constancy, mapping a constantly changing stream of retinotopic input onto an internal world-centered representation: a process referred to as ‘spatial updating’ (11). In principle, from a neurobiological perspective, there are indications that spatial updating in the brain can be exploited by visual prosthetics in a subpopulation of blind individuals who have intact eyes and residual gaze control. However, this might require active compensation with eye tracking and currently there is scarce literature that systematically compares the relative benefits of such compensation strategies.

When considering the biological requirements, it is relevant to note many individuals with late-acquired blindness have intact eyes and some level of gaze control (12, 13). In many cases of vision loss-related oculomotor difficulties the saccadic accuracy can be improved through training (14, 15), as the oculomotor system is under constant adaptive visual control (16). Similarly, it is speculated that improved oculomotor control can be regained with the reintroduction of visual feedback through phosphene vision. This suggestion is endorsed by exploratory clinical research. For instance, users of the Alpha IMS retinal photodiode implant (Retina Implant AG, Reutlingen, Germany) are found to display adequate scanning eye movements and object localization with little training and soon after implantation (17). Although these results may not directly translate to other types of prostheses, they suggest that spatial updating in the brain can be successfully exploited to enable natural-like visual scanning behaviour in visual prosthetics.

A crucial issue, however, is that visual scanning and spatial updating in the brain work under the assumption that the visual input changes when making eye-movements, but in most contemporary visual prostheses this is not the case. Unlike implants with retinal sensors, most prosthetic designs include a head-mounted camera that samples head-steered, rather than gaze-steered visual input. This has two interrelated implications: 1) In head-steered prostheses, only head movements will be available to scan the visual environment, while eye movements do not change the visual input. 2)

Phosphenes are retinotopically encoded, and they are coupled to eye movements (18–20). Therefore the same visual input will be perceived at a different location, depending on the gaze direction. An illustration of this ‘gaze-locked’ phosphene percept that is produced by head-steered prostheses is displayed in Figure 1a. Gaze-locked phosphene perception has been well-characterized in the prior literature (18– 23), and while it is known to cause localization problems (21), it is unclear to which extent these will restrict users in their daily life activities.

**Fig. 1.**
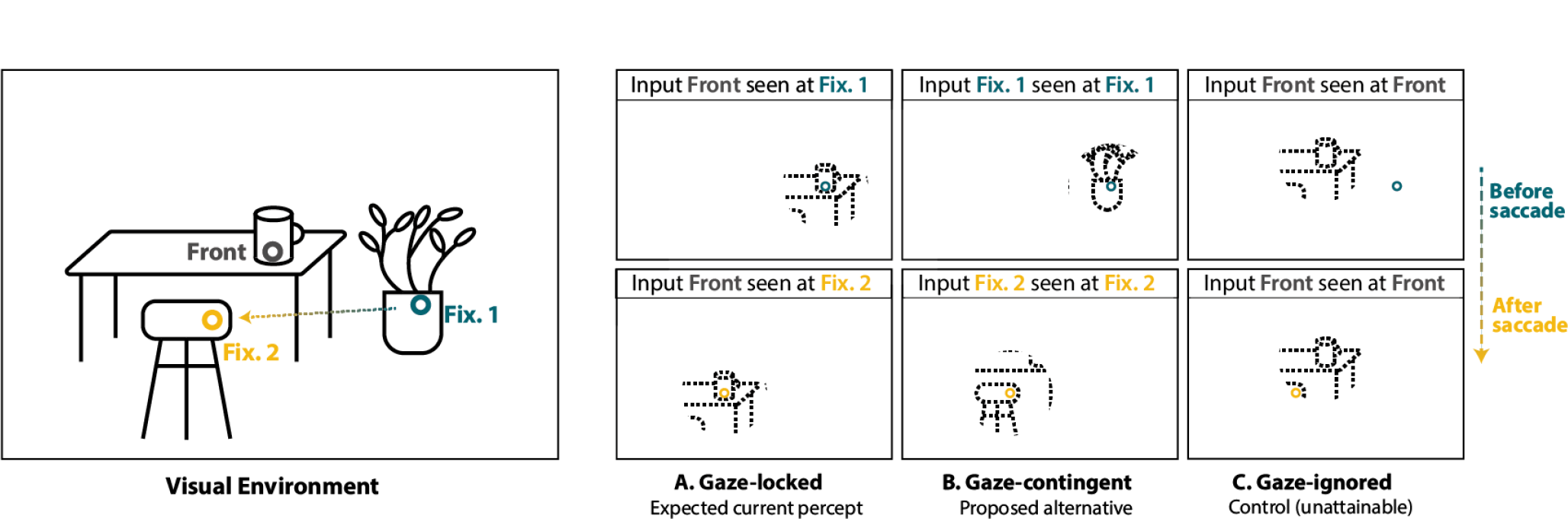
Schematic illustration of the three different simulation conditions used in our study. Left: illustration of the visual environment. Right: illustration of the simulated phosphene percepts before (top row) and after a saccade (bottom row). A) The gaze-locked condition simulates the perceptual experience of a prosthesis user with a head-steered visual prosthesis without eye-tracking compensation: the phosphenes encode what is straight in front of the head-mounted camera, but the phosphene locations are coupled to eye-movements. B) The gaze-contingent simulation condition emulates the percept created by a visual prosthesis with gaze-contingent eye-tracking compensation. The phosphene locations are gaze-centered (similar to the gaze-locked condition), but instead of head-centered visual input, the phosphene patterns encode a specific region of interest that matches with the gaze direction. C) The gaze-ignored simulation neglects any gaze-dependent effects on phosphene vision and it serves as a control condition which is unattainable in reality. In the gaze-ignored simulation, the phosphenes encode head-steered visual information (similar to the gaze-locked condition), but the phosphene simulation is displayed at a stationary location in the center of the head mounted display.

To mitigate the disorientation of gaze-locked phosphene percepts in head-steered prostheses, users typically undergo training to suppress eye movements, but so far results are disappointing as localization problems are reported to persist even years after implantation (21). Recently, the incorporation of a compensatory eye-tracking system with gaze-contingent image processing has emerged as a potential solution (22, 24–27). As illustrated in Figure 1b, gaze-contingent processing mimics natural visual sampling by processing only a gaze-centered region of interest (ROI) from a widefield camera stream: the gaze determines which region of the camera input is used to generate the phosphenes, such that the phosphenes display information about the specific location in the environment that is targeted by the gaze direction.

Several prior studies have investigated the benefits of gaze-contingent image processing. For instance, Caspi et al. (22) performed a target localization task on a computer screen with users of the Argus II retinal implant (Second Sight Medical Products, CA, USA) and found that gaze-contingent image processing resulted in higher precision and less head movements. Another study by Paraskevoudi and Pezaris (25) used a simulated prosthetic vision (SPV) paradigm with sighted subjects to non-invasively evaluate the functional quality of the artificial percept in a reading task. The authors found that gaze-contingent processing drastically improved the reading speed and accuracy compared to the control condition with head-steered vision.

These results are encouraging, but since these studies were performed in relatively controlled settings (for instance, participants sit behind a computer screen to identify target objects or letters), it remains unclear how these results translate to more natural settings and more complex activities. Given the opportunities for more immersive simulations to test more intricate tasks such as mobility and orientation (for example, see (28, 29)), it is relevant to explore the benefits of gaze-contingent processing for these tasks.

The current study aims to investigate gaze-contingent image processing in complex mobility and orientation tasks, using a mobile eye tracker in a virtual reality (VR) simulation. Specifically, we compare gaze-contingent image processing with a gaze-locked simulation of prosthetic vision, and a second control condition with gaze-ignored head-steered vision. We hypothesize that, owing to the availability of gaze-steered visual scanning and reestablished spatial updating, gaze-contingent phosphene vision yields higher mobility and orientation performance compared to gaze-locked and gaze-ignored phosphene vision. Note that the current study focuses on cortical prosthetic vision in particular, which, like natural vision, is characterized by an inhomogeneous resolution of information across the visual field, due to cortical magnification. Ultimately the goal of the current work is to provide further insights in the functional improvements that can be gained with an eye tracking system for gaze-contingent processing in head-steered cortical visual prostheses.

## 2. Materials and Methods

We performed two separate SPV experiments in indoor VR environments. A general description of the materials and methods used in our study and the specific details of Experiment 1 (obstacle avoidance) and Experiment 2 (scene recognition and visual search) are explained in the following sections.

### 2.1. Participants

In total, we recruited 43 healthy adult participants (23 for Experiment 1, and 20 for Experiment 2), with normal or corrected to normal vision, no limiting mobility impairments and no relevant prior history of cybersickness / motion sickness. None of the participants were familiar with the experimental tasks. One of the participants in Experiment 2 experienced VR-induced nausea during the practice session, and had to be excluded from the experiment, leaving a total of 19 subjects that completed Experiment 2. A descriptive summary of the study populations in both experiments can be found in Table 1. All participants provided written informed consent to participate in this study. This study was performed in accordance with the Declaration of Helsinki and the Nurenberg Code and was approved by the local ethical committee (ECSW, Radboud University).

**Table 1.** Average age and height (± standard deviation) of the study participants.

|  | Exp. 1 | Exp. 2 |
| --- | --- | --- |
| n | 23 | 19 |
| Age | 24.3 ($\pm 1.9$) | 24.1 ($\pm 2.9$) |
| Height | 173.4 ($\pm 8.0$) | 171.4 ($\pm 14.3$) |

### 2.2. Materials

For the VR simulation, we used the HTC VIVE Eye Pro (HTC Corporation, Taiwan) head mounted display (HMD), which has an inbuilt eye-tracker manufactured by Tobii eye tracking systems (Tobii AB, Sweden). The head and body position were tracked using the external cameras (base stations) provided with the VR system. The HMD was connected to a laptop (Dell Precision 7550 with NVIDIA Quadro RTX 4000 GPU) by wire. All wires were suspended from the ceiling to enable free movement.

### 2.3. Phosphene Simulation

For the VR simulation of cortical prosthetic vision, we used the Unity game engine (Unity Technologies, CA, USA) with different virtual 3D environments. For the real time integration of sensor data (subject pose, and gaze direction), we used the SDK (SRanipal) provided by the manufacturer. The simulation of cortical prosthetic vision was based on (30), adapted for Unity using the Cg shader programming language for graphics processing. To restrict the experimental complexity, we did not include temporal dynamics in the phosphene simulation, and we assumed full control over the phosphene brightness (with a high dynamic range). In the first experiment, we used 700 possible phosphene locations (i.e. simulating 700 electrodes), covering a field of view of 35 degrees. In the second experiment the number of phosphenes was reduced to 500 phosphenes to measure in minimal-vision conditions. The phosphene counts were determined using pilot experiments and are chosen for their associated level of difficulty: with the current values, the experimental tasks were difficult, but could still be performed. The visual field coverage was purposely limited to a small area to reflect possible limitations to the area that can be used for the electrode implantation. Furthermore, we expected the effects of the different gaze conditions to be more pronounced when phosphenes cover only a relatively small field of view. The encoding of the phosphenes (i.e. the activation pattern) was based on scene simplification with surface normal estimation combined with edge detection, which, also based on pilot experiments, turned out to be an intuitive phosphene representation for summarizing the current visual environments.

### 2.4. Experimental conditions

Our study compares three experimental conditions (see Figure 1). In the gaze-locked simulation condition, the phosphenes encode what is captured by the head-centered virtual camera and the phosphene locations are coupled to eye movements. Note that while eye-tracking is required to simulate gaze-locked phosphene vision, the study condition simulates the experience produced by a head-steered visual prosthesis *without* an eye-tracking system. The gaze-contingent simulation condition simulates the effect of including a compensatory eye-tracking system in a head-steered prosthesis. Here, the phosphenes encode a region of the visual environment that matches with the gaze direction, enabling the user to sample visual input with eyemovements as well as head movements. The third study condition, the gaze-ignored condition, resembles commonly-used phosphene simulations that ignore the effects of eye movements on phosphene perception. Note that this simulation condition is included as a control condition to measure the isolated effects of head-steered visual sampling, and it does not accurately reflect the perceptual experience of cortical visual prosthesis users.

### 2.5. Experiment 1: Obstacle Avoidance

#### 2.5.1. Virtual Environment

The design of the virtual environment was based on a previous (real-world) mobility study (29), and featured a straight corridor with boxes that served as obstacles (see Figure 2b and Figure 4). The virtual corridor was 44 meters long and consisted two empty sections and a middle section that contained 19 sets of obstacles. The obstacles were spaced evenly along the length of the corridor, two meters apart. The obstacle sets consisted either of a single small box or a combination of two large boxes, that were arranged similar to a partial wall or door-frame, spanning the entire height of the corridor and blocking two-thirds of its width. The placement of the obstacles varied between trials, but always matched one of the three unique templates. In total, each of the three hallway variants with a unique obstacle placement was always visited three times by each participant (once for every condition).

**Fig. 2.**
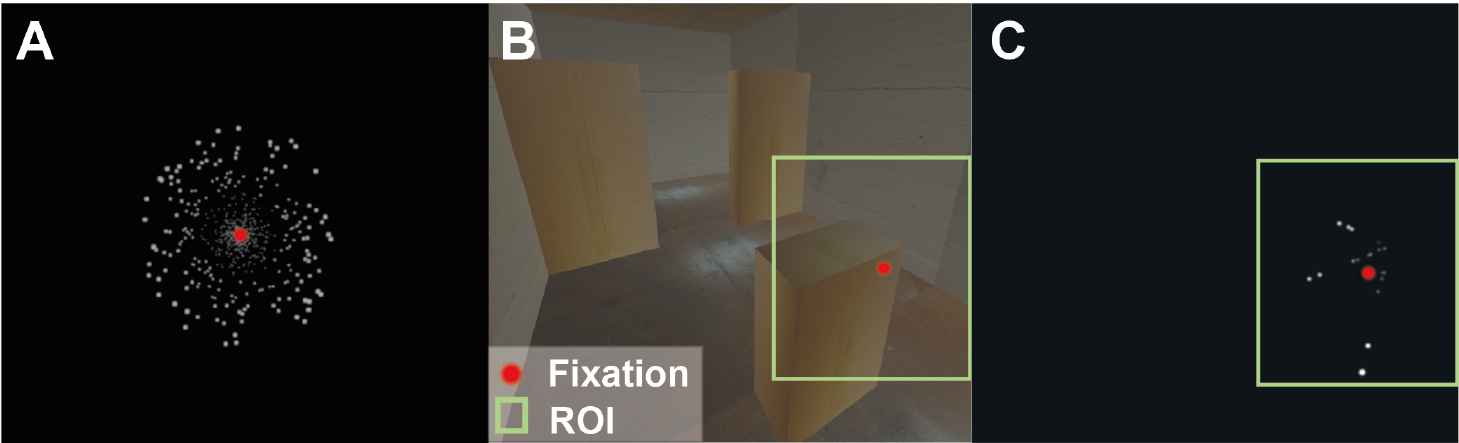
The phosphene simulation that was used in Experiment 1. A) Simultaneous activation of all 700 possible phosphenes centered at the point of fixation. B) The virtual hallway environment that was used in the practice session of Experiment 1. The green rectangle indicates the region of interest that would be used for gaze-contingent image processing, if the participant looks at the fixation point indicated in red. C) The resulting phosphene simulation after processing the region of interest in panel B. Note that only the phosphenes are activated that are located on the surface boundaries.

**Fig. 3.**
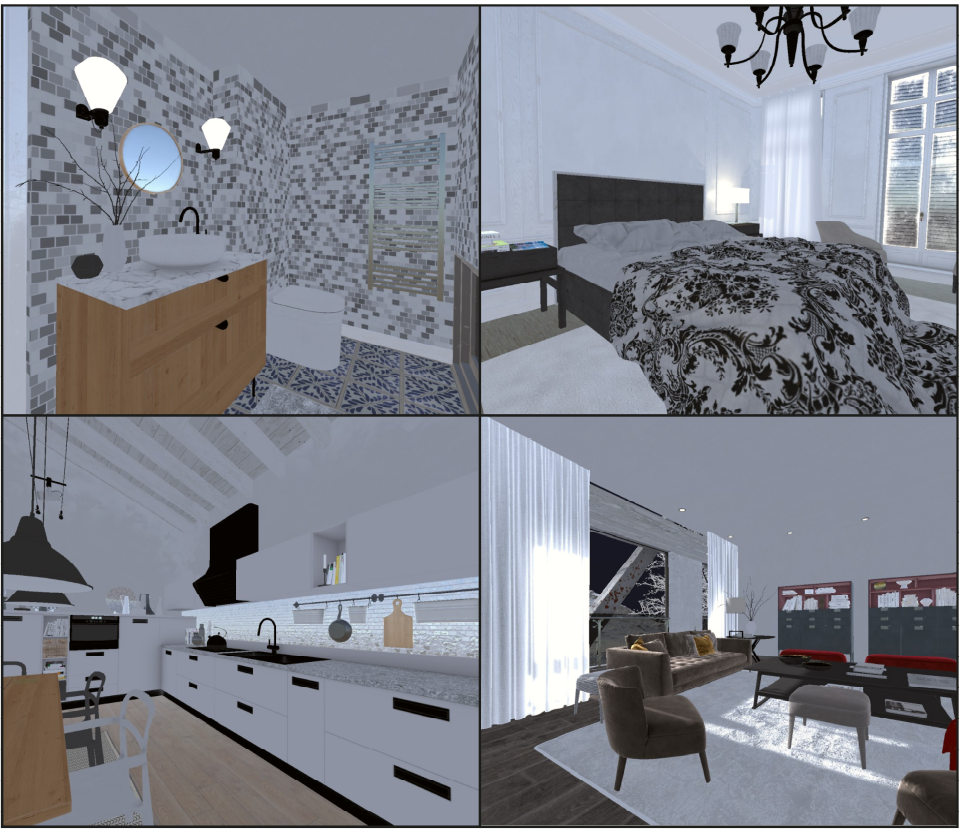
Example virtual indoor environments that were used for the scene recognition task in Experiment 2. The four possible room categories were: bathroom, bedroom, kitchen and living room.

**Fig. 4.**
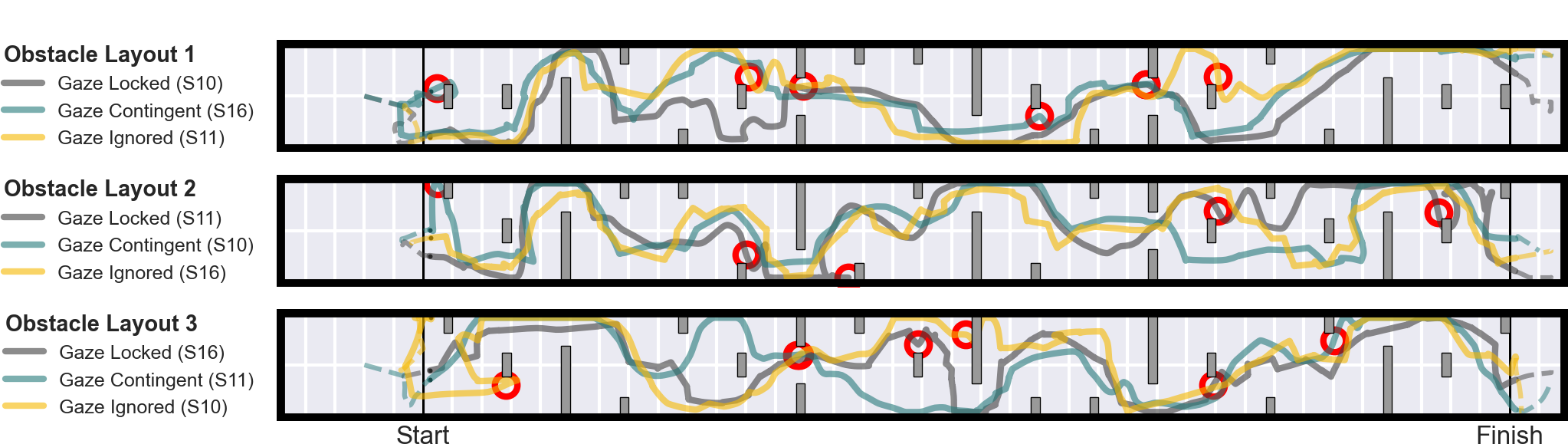
Hallway layouts and mobility trajectories in Experiment 1. The virtual hallway was 44 meters long, and contained 19 obstacle arrangements that were spaced evenly across the length of the hallway, two meters apart. The number of collisions and the trial duration was recorded between the startand finish line (37 meters distance). Each row displays one of the three unique variations of obstacle placements. The head position in the virtual hallway is plotted of three representative subjects (S10, S11 and S16). In total, nine trials are visualized. The obstacles are displayed as dark rectangles. Collisions with obstacles are indicated with a red circle.

#### 2.5.2. Movement Control

While using the VR equipment the participants were standing still in a spacious room without hazards. The participants were instructed to keep their feet in place, but are free to look around and use their hands to explore the environment. They moved through the virtual hallways using the circular touch pad of the VR handheld game-controllers. By placing their thumb on the touch pad, the participants could control their speed and direction in the virtual environment, allowing for a maximum speed of 2.0 ms^*−*1^. Upwards on the trackpad corresponded to the direction participants were looking at, so that “forward” on the trackpad would be aligned with what participants perceive as “forward”. In case the participants collided with an obstacle in the virtual environment, they received haptic feedback through a short and strong vibration signal of the VR controller. The participants could also use the VR controllers to explore the edges of the obstacles: in case their virtual hand position in the hallway intersected with an obstacle, the controllers pulsed with a weaker vibration to inform the participant that they were touching the obstacle.

#### 2.5.3. Training session

All participants performed the experiment in a single one-hour visit, which consisted of a two-part training period, the actual data collection, and a final questionnaire. In the training period, participants were placed in a short corridor featuring two example obstacles. First, the participants practiced moving in the VR environment using the game-controllers. The participants practiced with natural vision (without phosphene vision), until they felt secure in controlling their movement to get past the training obstacles and understood the different vibrations of the collision feedback. Next, a contour filter was applied, reducing the visual fidelity, and more training was performed until the participants securely navigated through the environment. Finally, the participants practiced navigating past the obstacles using the phosphene simulation. An even amount of training was performed with the three different phosphene simulation conditions. The participants could switch back and forth between conditions until they were able to safely navigate the test corridor.

#### 2.5.4. Experimental session

The experiment was split into three blocks of three trials each. In each block participants would see each variation of object placement and each condition exactly once. The order of the conditions and object layouts inside each block were randomized for each participant. After completing all three blocks, participants had performed all nine possible combinations of the simulation conditions and the variants in obstacle placement. After each block and at the end of the experiment, the participants filled out a survey to collect information on the subjective experience.

#### 2.5.5. Experimental task

Participants were instructed to avoid collisions as well as they can while still moving as fast as possible. The participants were not informed about the simulation condition before starting each trial. During the experiment, participants kept their feet in place and were encouraged to survey and explore their environment with their hands and by looking around.

### 2.6. Experiment 2

With a different set of participants, the second experiment evaluated visual orientation using a scene recognition task and a visual search task. Both tasks were tested in separate trials. In the scene recognition trials, subjects were placed in a randomly selected unfamiliar virtual environment in which they had to identify the room category (bathroom, bedroom, kitchen or living room). In the visual search trials, subjects were situated in a familiarized studio environment, in which they were tasked to sequentially find many prompted target objects within a given time limit of 2 minutes.

#### 2.6.1. Virtual environments

We used several virtual environments designed by ArchVizPro Studio (Oneiros srl, Italy), obtained from the Unity asset store (Figure 3. The environments were slightly modified for the experiments: some rooms and objects were re-scaled and some objects were removed or added to create a more realistic and representative representation. In total we used 16 different environments. One environment, the studio environment that was used for the visual search task, was visited frequently to allow familiarization by the subjects. The other 15 environments (three bathrooms, four bedrooms, three kitchens and five living rooms) served as the unfamiliar environments for the scene recognition tasks. The unfamiliar environments were visited maximally twice by the same subject and were only observed with the simulated phosphene vision. The familiarized studio environment was also observed with natural sight (i.e. with deactivated phosphene simulation) during the practice session at the beginning of the experiment.

#### 2.6.2. Movement Control

In contrast to Experiment 1, the camera pose in the virtual environments was entirely dependent on the participant’s pose in reality (the position of the headset), and movement control of the character by using the trackpad on the VR game-controller was disabled. Subjects were instructed to stand in the middle of the room and to look around in all directions. Subjects could turn around in place, but were instructed to avoid walking away from the start position. Between trials, subjects were redirected to the start location and orientation if necessary, facing an instruction screen in the VR environment.

#### 2.6.3. Practice Session

The total experiment lasted maximally two hours, starting with a practice session of roughly 30 minutes. In the practice session, the subjects could get adjusted to the VR experience, the phosphene simulation and the experimental tasks. The practice session included several trials with free exploration, a visual search trial and three scene recognition trials. The environments that were used for practicing the scene recognition task were never reused in the actual scene recognition session.

#### 2.6.4. Scene Recognition Session

After the practice session, subjects performed a scene recognition session that lasted roughly one hour, including breaks. The scene recognition session consisted of three blocks, one for each condition, plus another repetition of three blocks. Each block contained three trials, each with a different environment. The instructions were to report the room category as fast as possible, while considering that accuracy is preferred over speed. To ensure that the total duration of the experiment did not exceed two hours, all scene recognition trials were limited by a maximal duration of two minutes, after which subjects were instructed to provide a (forced) choice of room category, even if they were not confident of their answer. The environments were randomly selected and the order of the experimental conditions were randomly shuffled to avoid systematic biases owing to environment characteristics, practice effects or fatigue. Before each scene recognition block, we included an additional practice trial with a visual search task in the studio environment. This served two purposes: 1) each time when the simulation condition was switched, the subject had the opportunity to practice and get adjusted to the new simulation condition before starting the scene recognition trial 2) by performing extra practice trials in the studio environment, during the scene recognition session, subjects became increasingly familiarized with the environment before the visual search session.

#### 2.6.5. Visual Search Session

Roughly the final 30 minutes of the experiment (including breaks) were reserved for the visual search session in the studio environment. By this time, subjects have visited the studio environment at least ten times for a duration of 1,5 minutes per visit. The visual search session consisted of three repetitions of three trials. Each trial lasted two minutes and tested one of the three study conditions. The task was to sequentially search and find as many target objects as possible (either: a pair of shoes, a mug or a stool). At every point in time, there was only one target and each target was only made visible when prompted. Once the participant has found the object, they were requested to indicate its location by looking at it, and pointing at it with the controller while pressing the response button. We used a preset of 14 possible locations where the target could appear and targets appeared at sensible locations To avoid targets from appearing in sight, the new target location was never close to the previous target location.

### 2.7. Study Outcomes

#### 2.7.1. Primary study outcomes

Several primary outcomes were used to compare the performance across study conditions: For the mobility task in Experiment 1, we analyzed the average trial duration and number of collisions per trial. For the scene recognition task, we measured the classification accuracy and the average trial duration. For the visual search tasks we analyzed the average search duration. Furthermore, in Experiment 2, we recorded a subjective evaluation metric after each trial.

#### 2.7.2. Additional outcomes

To evaluate behavioral strategies and differences in visual exploration, we inspected the eye-gaze trajectories in Experiment 2, and calculated the angular velocity of eye and head rotations. Furthermore, we analyzed the questionnaires which were filled out at the end of each block in Experiment 1 and at the end of Experiment 2 to record self-reported observations.

### 2.8. Data Analysis

#### 2.8.1. Pre-processing

The data obtained in both experiments were analyzed in Python (version 3.11.0). In Experiment 1, five trials were excluded from the analysis because of missing data (a portion of the frames was not saved successfully). To obtain the average number of collisions per trial, the head position was projected on the horizontal plane, and frames were marked as a collision in case the distance between the projected head position and the closest obstacle was smaller than 0.225 meters. The trial duration in the mobility task was calculated as the time between crossing the start and finish lines, which were separated 37 meters.

In Experiment 2, in the scene recognition task, three subjects did not perform better than chance level across all conditions on average. These subjects were excluded from the analysis. For the visual search task, the gaze directions were analyzed to verify whether subjects found the correct target. Target reports that deviated more than 15 degrees from the correct target direction were considered invalid. Two subjects reported the wrong target object in more than 40% of the cases and were excluded from the analysis. Two different subject did not find a minimum of three targets per study condition and were excluded from the analysis. One target object was not correctly found in a minimum of 40% of the cases and was therefore excluded from the analysis as well. Only search durations of successful searches (targets that were correctly found) were analyzed. We verified that the exclusion of data did not alter the conclusions of the analysis.

#### 2.8.2. Statistical analysis

For each subject we averaged the endpoints across trials. A Shapiro-Wilk test was used to test if data was normally distributed. To test for significant differences between study conditions, we used a paired samples t-test (for normally distributed endpoints) or a Wilcoxon signed-rank test (for non-normally distributed endpoints). We used a Bonferroni correction to correct for the number of comparisons (three for each endpoint), rendering a required significance level of α = 0.0167. To explore potential learning effects over the course of the experiment, we performed a linear regression analysis, also testing for possible interaction effects between the study conditions.

## 3. Results

### 3.1. Primary outcomes

#### 3.1.1. Exp. 1 Obstacle avoidance

Several example mobility trajectories are displayed in Figure 4. The primary results of the analysis of Experiment 1 are displayed in Figure 5. The average number of collisions was significantly higher in the gaze-locked study condition compared to the gaze-contingent condition (p = 0.014) and the gaze-ignored condition (p *<* 0.001). Furthermore, the average trial duration was significant higher in the gaze-locked study condition compared to the gaze-contingent study condition (p *<* 0.001) and the gaze-ignored study condition (p *<* 0.001). The regression analysis revealed no significant reduction in the number of collisions across subsequent blocks, but there was a significant reduction of the trial durations (14% per block; p *<* 0.001). The slopes for the different study conditions were not significantly different, indicating no interaction effects.

**Fig. 5.**
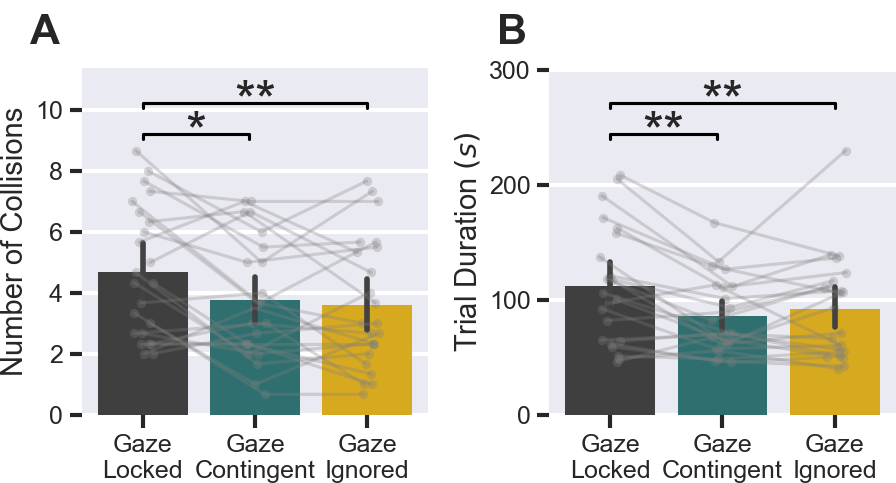
Primary results of the mobility task in Experiment 1. A) Average number of collisions per trial (N=23, Paired-samples t-test). B) Average trial duration in seconds (N=23, Wilcoxon signed-rank test). ^*^p *<* 0.0167; ^**^p *<* 0.003; ^***^p *<* 0.0003.

#### 3.1.2. Exp. 2 - Scene recognition

The results of the scene recognition task are displayed in Figure 6 (panels a-c). The scene recognition accuracy was not significantly different in the gaze-contingent vision condition compared to the gaze-locked condition, but gaze-contingent vision yielded significantly lower trial durations (51 versus 63 seconds on average, p = 0.0012). Furthermore, the subjective rating was significantly higher, by more than 1 point on average (on a scale of 1 to 7; p *<* 0.001). The gazeignored control condition was characterized by significantly higher scene classification accuracy compared to the gazecontingent condition and the gaze-locked condition (p = 0.0067 and p = 0.012, respectively), as well as higher subjective ratings (p = 0.016 and p *<* 0.001). The trial durations in the gaze-ignored condition were significantly shorter compared to the gaze-locked condition (p *<* 0.001), but not compared to the gaze-contingent vision condition (p = 0.061).

**Fig. 6.**
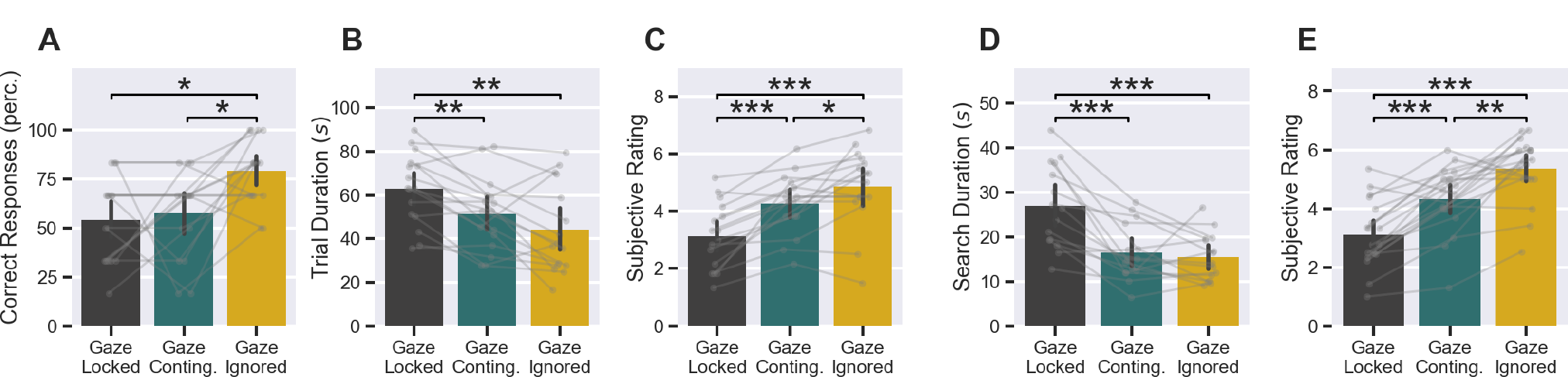
Primary results of the scene recognition task (A-C) and the visual search task (D, E) in Experiment 2. A) Classification accuracy in the scene recognition task (N=16). B) Average trial duration in the scene recognition task (N=16). C) Average reported subjective rating in the scene recognition task (N=16). D) Average search duration in the visual search task (N=15). E) Average reported subjective rating in the visual search task (N=15). ^*^p *<* 0.0167; ^**^p *<* 0.003; ^***^p *<* 0.0003. (panels A and E: Wilcoxon signed-rank test; panels B,C and D: Paired-samples t-test) .

#### 3.1.3. Exp 2 Visual search

The results of the visual search task are displayed in Figure 6 (panels d and e). Search durations were significantly lower in the gaze-contingent processing condition compared to the gaze-locked control condition (27 versus 16 seconds on average, respectively, p *<* 0.001), and the subjective ratings were significantly higher by more than 1 point (p *<* 0.001). No significant difference in search duration was found between the gaze-ignored and the gaze-contingent condition. However, the gaze-ignored control condition was characterized by significantly lower search durations compared to the gaze-locked condition (p *<* 0.001). Furthermore, subjective ratings were significantly higher in the gaze-ignored condition compared to the gaze-contingent and gaze-locked study conditions(p = 0.0017 and p *<* 0.001, respectively).

### 3.2. Secondary outcomes

#### 3.2.1. Gaze analysis

Figure 7 displays several example gaze trajectories from Experiment 2. In the visualized trials it can be observed that the gaze-contingent and the gaze-locked study conditions were characterized by a larger spread compared to gaze-ignored vision. Averaged over all participants, this effect was found to be significant (p *<* 0.001 for both comparisons). The results of analyzing the angular eye- and head velocities are displayed in Figure 8. The gaze velocity in the gaze-contingent study condition was higher compared to the gaze-locked condition (p *<* 0.001), but lower compared to the gaze-ignored condition (p *<* 0.001). Also, the head velocity was significantly higher in the gaze-contingent study condition compared to the gaze-locked study condition (p *<* 0.001), but lower compared to the gaze-ignored study condition (p = 0.0062). Compared to the gaze-ignored study condition, the gaze-locked condition was characterized by a lower gaze-velocity (p *<* 0.001) and a lower head velocity (p *<* 0.001).

**Fig. 7.**
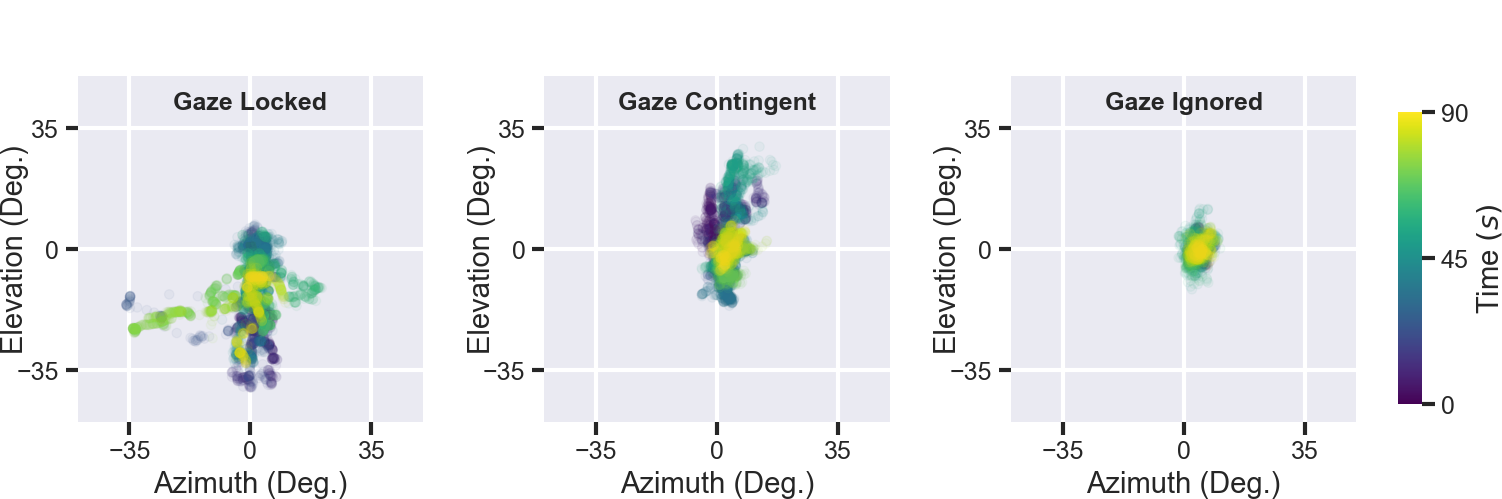
Example eye-gaze trajectories (raw data) in three trials of a representative participant in Experiment 2. Colors indicate the elapsed time. Each of the visualized trials had a different study condition (from left to right: gaze-locked, gaze-contingent, and gaze-ignored).

**Fig. 8.**
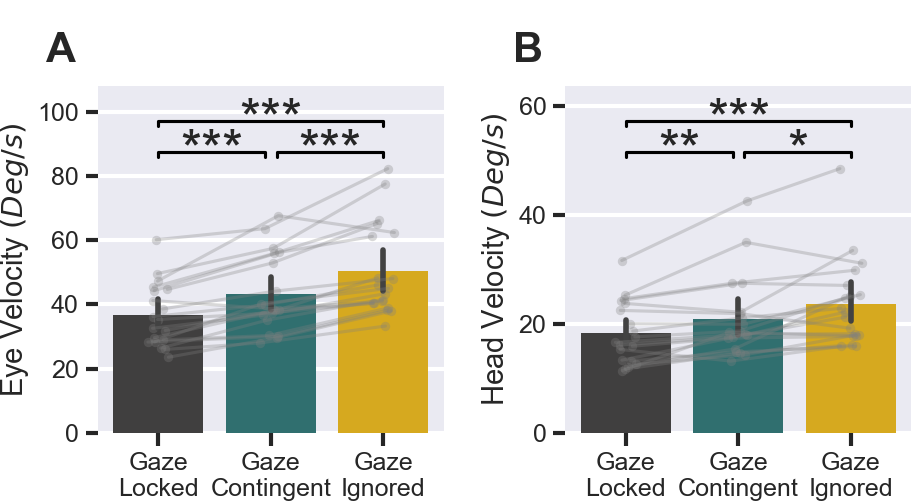
Analysis of the angular eye- and head velocity in Experiment 2. A) Average eye velocity. B) Average head velocity. ^*^p *<* 0.0167; ^**^p *<* 0.003; ^***^p *<* 0.0003 (N=19, Wilcoxon signed-rank test).

#### 3.2.2. Questionnaires

After each block in Experiment 1, the participants ranked the subjective experience of the trials in that block. In 68% of the responses, trials with the gaze-locked condition were ranked as the most tiring, versus 13% and 19% of the responses, respectively, ranking the gaze-contingent trial or the gaze-ignored trial as the most tiring. In 77% of the responses, the trial with the gaze-locked condition was marked as the least comfortable versus 9% and 14%, respectively, ranking the gaze-contingent or the gaze-ignored trial as the least comfortable. In terms of subjective functionality (“How well can you navigate with this condition?”) the gaze-contingent vision trial was ranked the highest in 48% of the responses, versus 14% for the gaze-locked condition and 38% for the gaze-ignored condition. In 27% of the responses, the participant indicated to experience motion sickness in the trials with gaze-locked vision, versus 6% for both the gaze-contingent and gaze-ignored vision. When asked about specific strategies, 57% of the participants reported that they used exaggerated head movements, 40% of the participants reported that they estimated depth by moving sideways and 61% of the subjects indicated that they estimated the object boundaries by scanning with the center-most phosphenes.

After completing Experiment 2, 37% of the participants reported that they had felt disoriented in the gaze-locked study condition, versus 11% in the gaze-contingent condition and 0% in the gaze-ignored condition. 11% of the subjects experienced nausea in the gaze-locked condition, 5% in the gaze-contingent condition and 0% in the gaze-ignored condition. 10% of the subjects report that they experience motion sickness regularly in everyday life. 68% of the subjects reported that they purposely tried to keep their eye still in some of the trials. 47% percent of the subjects reported that despite trying they found it difficult to suppress their eye-movements. 74% of the subjects reported that their eyemovement strategy depended on the eye-tracking condition. When asked about other specific strategies throughout the second experiment, 53% of the subjects indicated that they used exaggerated or frequent head movements, 21% of the subjects indicated that they estimated depth by moving sideways, 47% of the subjects indicated that they used the centermost phosphenes to estimate object boundaries.

## 4. Discussion

In this simulated prosthetic vision study, we evaluated the functional benefits of gaze-contingent image processing with eye-tracking in several mobility and orientation tasks. Across both experiments, we found, in line with our hypothesis, a reduced number of collisions, lower trial durations and higher subjective ratings for the gaze-contingent study condition compared to the gaze-locked condition. These results suggest that the inclusion of a compensatory eye-tracking system in head steered visual prostheses may improve the functional quality of head-steered visual prostheses in mobility and orientation. Furthermore, in the second experiment, the gaze-ignored control condition generally outperformed both other simulation conditions in all tasks. We speculate that this may be partly owed to adverse effects of an imperfect eye-tracking system. In addition, our findings point to the relevance of considering the effects of gaze in phosphene simulation studies, as neglecting gaze can produce overoptimistic outcomes.

### 4.1. Benefits of gaze contingency with eye tracking

There was no significant improvement of the classification accuracy in the scene recognition task with gaze-contingent versus gaze-locked phosphene vision. However the lower number of collisions in the mobility task and the higher speed and subjective ratings across the various tasks in both experiments are in line with the results of previous work (22, 24– 27, 31, 32). Several prior studies have specifically compared gaze-contingent with gaze-locked vision in visually-guided tasks, such as searching letters (24), pointing to targets on a screen (22, 26) or even reading (25). In addition to the aforementioned studies, that found an overall improvement with gaze-contingent image processing, the current results suggest that the improvements with gaze-contingent image processing extend to more complex situations such as 3D orientation and mobility with high phosphene counts. These findings underscore the importance of gaze-compensation for gaze-locked vision in head-steered prostheses. For the clinical cases in which eye-tracking is feasible, gaze-contingent image processing is expected to yield universal advantages for the functional and subjective quality of the prosthetic vision.

### 4.2. Superior performance in gaze-ignored simulation

In Experiment 1 we found no significant difference in performance between gaze-contingent and gaze-ignored vision. The lower overall performance (i.e. lower scene recognition accuracy and lower subjective ratings) with gaze-contingent phosphene vision in Experiment 2 compared to gaze-ignored vision was unexpected. These findings are in contrast with results of a prior study by Titchener et al. (26), who found that gaze-contingent simulations yielded similar or even better target localization performance compared to gaze-ignored vision. Although we cannot be certain, we speculate that inaccuracies in our mobile eye-tracking system may have caused adverse effects on the performance (see Section 4.6). Imperfect gaze estimations may have caused discomfort or perceptual disturbances, masking potential benefits of the gaze-contingent phosphene simulation.

In any case, the relatively high performance with the gaze-ignored control condition support the observation that adequate functionality can be achieved with head-steered visual scanning only. In natural vision, actions are strongly directed and guided by eye-movements (33). Nevertheless, our results are in line with in prior clinical work (34) and simulated prosthetic vision studies (e.g. (32)), showing that a lack of gaze-assisted visual scanning is surmountable. In this light, the functional limitations of gaze-locked artificial vision in head-steered prostheses (such as described by Sabbah et al. (21)) likely do not originate from the restricted visual scanning, but from conflicts in spatial updating.

### 4.3. Implications of neglecting gaze in simulations

Although the gaze-locked perception of phosphenes in head-steered prostheses has been well-characterized in the prior literature (18–23), eye movements are commonly ignored in simulation studies. Our finding that the performance with gaze-locked phosphene vision is significantly lower than with gaze-ignored phosphene vision is in line with results from prior work (23) and this may hold even after years of training (21). Since simulations that neglect eye-movements evaluate a condition that is unattainable in the clinical setting, it is important to consider that studies with gaze-ignored simulations may yield overoptimistic expectations of the functional performance with visual prostheses. For more realistic evaluation of functional value of prosthetic vision, it is therefore crucial to include eye-tracking in the experimental setup, similar to, for instance, (24–27, 31, 32).

### 4.4. Learning effects

Importantly, the use of prosthetic vision, and especially gaze-locked vision requires intensive training (21). In both experiments, most of the participants reported that they had sufficient training with the simulation conditions during the practice trials. Indeed, we did not find a significant training effect in the primary experimental outcomes, with one exception: the reduced trial duration in Experiment 1 indicated improved mobility performance over the course of the experiment. We found no interaction between the different study conditions. Although previous suggested that gaze-contingent viewing can be adopted with relatively minimal effort Paraskevoudi and Pezaris (25), we did not find any differences in the learning abilities with the different conditions.

### 4.5. Strategies

The participants did not receive any specific instructions for coping with the different gaze conditions. The reduced gaze-velocity in the gaze-locked compared to the gaze-contingent study condition indicates that subjects intuitively learned to suppress eye-movements with gaze-locked vision. Indeed, the majority of subjects indicated in the exit interview of Experiment 2 that they tried to keep their eyes still. Note that most of the subjects that tried to actively suppress eyemovements experienced difficulties in succeeding so. These responses are similar to the experiences described by users of head-steered prostheses (21, 22), suggesting fundamental challenges with compensation training based on eyemovement suppression. Some complicating factors are reflexive eye-movements (e.g. vestibulo-ocular reflex) which remain difficult to suppress. When asked about other strategies, more that half of the participants in both experiments indicated that they used frequent or exaggerated head movements. Interestingly, the average head velocity in Experiment 2 was significantly higher with gaze-contingent vision compared to the gaze-locked condition. This finding was unexpected, as the reestablished visual scanning with eye-movements would alleviate the need for head movements. We speculate that the reduced rotational head velocity with gaze-locked vision partly reflects uncertain scanning behaviour caused by perceptual instability. One participant indicated in the free written responses that they ‘decreased the exaggerated head movements due to nausea’. In both experiments, the side effects such as disorientation and nausea were reportedly lower with the gaze-contingent versus the gaze-locked vision. Noteworthy, there was a large variety in self-reported strategies for performing the experimental tasks. Incontestably, besides the functional performance, it is important to consider these subjective measures of convenience and (dis)comfort for the development of visual prostheses, accounting for personal differences in preference.

### 4.6. Limitations and future directions

There are several limitations to the current study. One practical limitation is the indirect user control in the mobility experiment that was used for moving through the virtual environment. The use of finger-steered movement with a sensitive trackpad may have led to difficulties interpreting the virtual self-motion. This holds especially for phosphene vision that lacks natural optical flow cues.

One limitation of the scene recognition task, was the limited generalizability of the 3D environments. Although there were large differences in the used scenes in terms of design and objects, the used environments reflect only a subset of the diversity (e.g. cultural diversity) of real-world environments. Due to the fundamental low-level visual differences between the experimental conditions, it is unlikely that the results are much influenced by the specific choice of environments.

There are several inherent limitations of simulated prosthetic vision with sighted subjects. Although efforts were undertaken to maximize the realism of the phosphene simulation, it nevertheless misses some of the true complexities of artificial phosphene vision, such as temporal dynamics, irregular shapes, and interactions between electrodes. Furthermore, blind individuals are likely to use different strategies than sighted study subjects (especially for mobility). Although we aimed to measure visually-guided mobility, the prospective users of visual prostheses may rely on multimodal sensory information (e.g. using a cane).

Lastly, and related, although the current study implemented an immersive simulation with complex 3D tasks, it is important to acknowledge that nevertheless there is a gap with the actual clinical situation. the tasks that were tested are still relatively controlled compared to the real life situation, where orientation and mobility is usually a closed-loop interaction with the environment. With the progression of prosthetic technology it will become possible to perform clinical experiments with more complex visually-guided activities of daily living. In general, further clinical work can improve the realism and relevance of simulation studies. Vice versa, simulation studies can give further directions for progressing the technology towards the clinical setting.

### 4.7. Conclusion

In this mixed reality simulated prosthetic vision study, we evaluated the benefits of eye tracking-based gaze compensation for head-steered prostheses in the context of mobility and orientation. We found that gaze-contingent processing improved the performance in all experimental tasks. Likely, this improvement is mostly owed to compensation for spatial updating conflicts and only to a lesser extent to extended freedom of visual scanning. We found that neglecting the effects of eye-movements in simulations of head-steered prostheses can yield overoptimistic results. This should be taken into account in future simulation work. Taken together, we conclude that for cases in which eye-tracking is feasible, gaze-contingent image processing is expected to improve the functional and subjective quality of head-steered prostheses.

## ACKNOWLEDGEMENTS

This work has received funding from the European Union’s Horizon 2020 research and innovation programme under grant agreement No 899287. This work was supported by the following grants of the Dutch Organization for Scientific Research (NWO): NESTOR (STW grant number P15-42), INTENSE (cross-over grant number 17619).

